# Substantial Neutralization Escape by the SARS-CoV-2 Omicron Variant BQ.1.1

**DOI:** 10.1101/2022.11.01.514722

**Authors:** Jessica Miller, Nicole P. Hachmann, Ai-ris Y. Collier, Ninaad Lasrado, Camille R. Mazurek, Robert C. Patio, Olivia Powers, Nehalee Surve, James Theiler, Bette Korber, Dan H. Barouch

**Author notes:** Corresponding author: Dan H. Barouch, M.D., Ph.D., Center for Virology and Vaccine Research, 330 Brookline Avenue, E/CLS-1043, Boston, MA 02115. **Authors for Print Edition** Jessica Miller, B.S., Nicole P. Hachmann, B.S., and Dan H. Barouch, M.D., Ph.D.; Center for Virology and Vaccine Research, Beth Israel Deaconess Medical Center, Boston, MA, USA.

## Abstract

Omicron BA.5 has been the globally dominant SARS-CoV-2 variant and has demonstrated substantial neutralization escape compared with prior variants. Additional Omicron variants have recently emerged, including BA.4.6, BF.7, BA.2.75.2, and BQ.1.1, all of which have the Spike R346T mutation. In particular, BQ.1.1 has rapidly increased in frequency, and BA.5 has recently declined to less than half of viruses in the United States. Our data demonstrate that BA.2.75.2 and BQ.1.1 escape NAbs induced by infection and vaccination more effectively than BA.5. BQ.1.1 NAb titers were lower than BA.5 NAb titers by a factor of 7 in two cohorts of individuals who received the monovalent or bivalent mRNA vaccine boosters. These findings provide the immunologic context for the rapid increase in BQ.1.1 prevalence in regions where BA.5 is dominant and have implications for both vaccine immunity and natural immunity.

---

Omicron BA.5 has been the globally dominant SARS-CoV-2 variant^1^ and has demonstrated substantial neutralization escape compared with prior variants^2,3^. Additional Omicron variants have recently emerged, including BA.4.6^4^, BF.7, BA.2.75.2, and BQ.1.1, all of which have the Spike R346T mutation (**Fig. 1A**). In particular, BQ.1.1 has rapidly increased in frequency (**Fig. S1**), and BA.5 has recently declined to less than half of viruses in the United States. The ability of BQ.1.1 to evade NAbs induced by vaccination and infection has not yet been determined.

**Figure 1.**
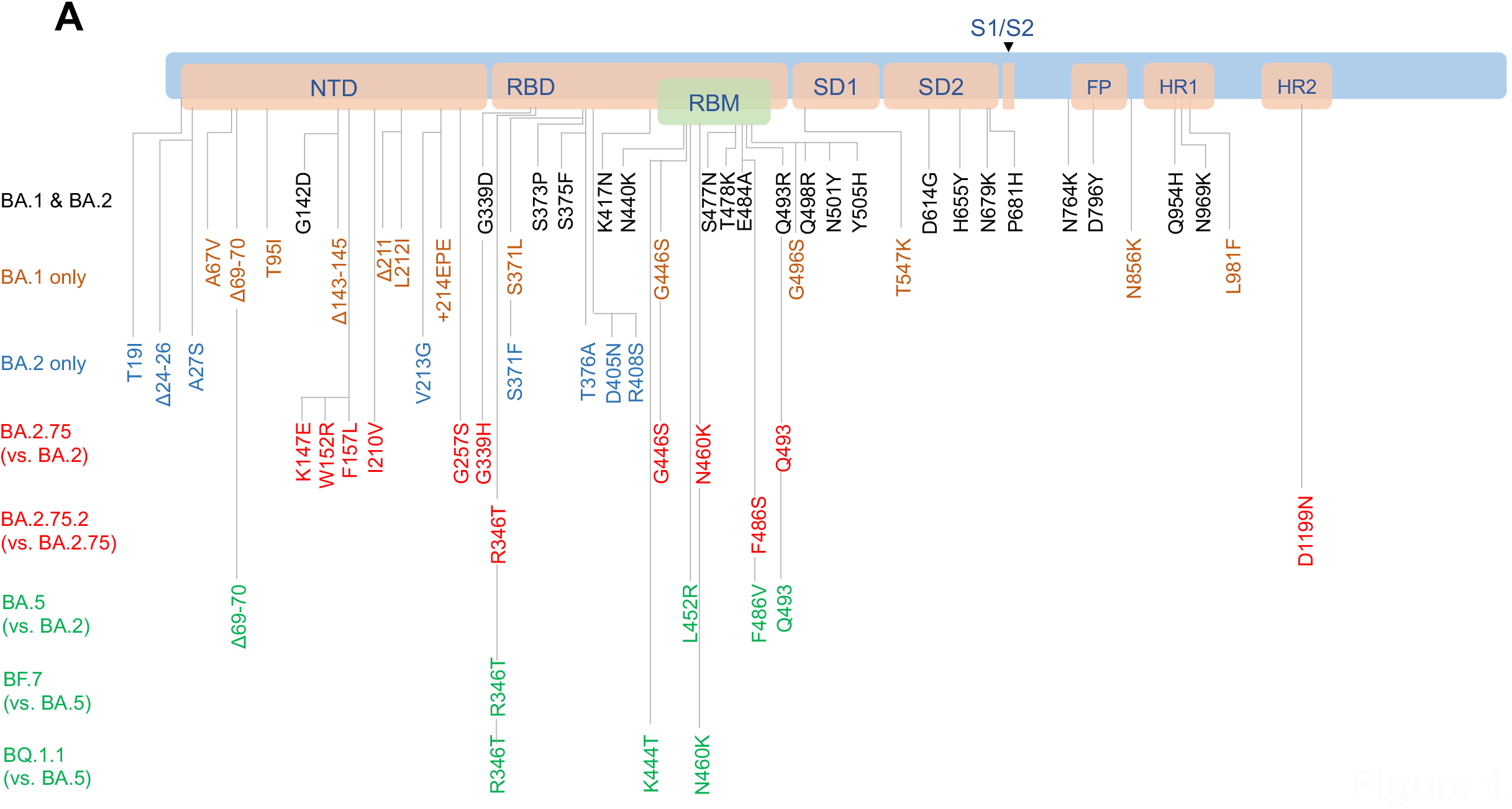
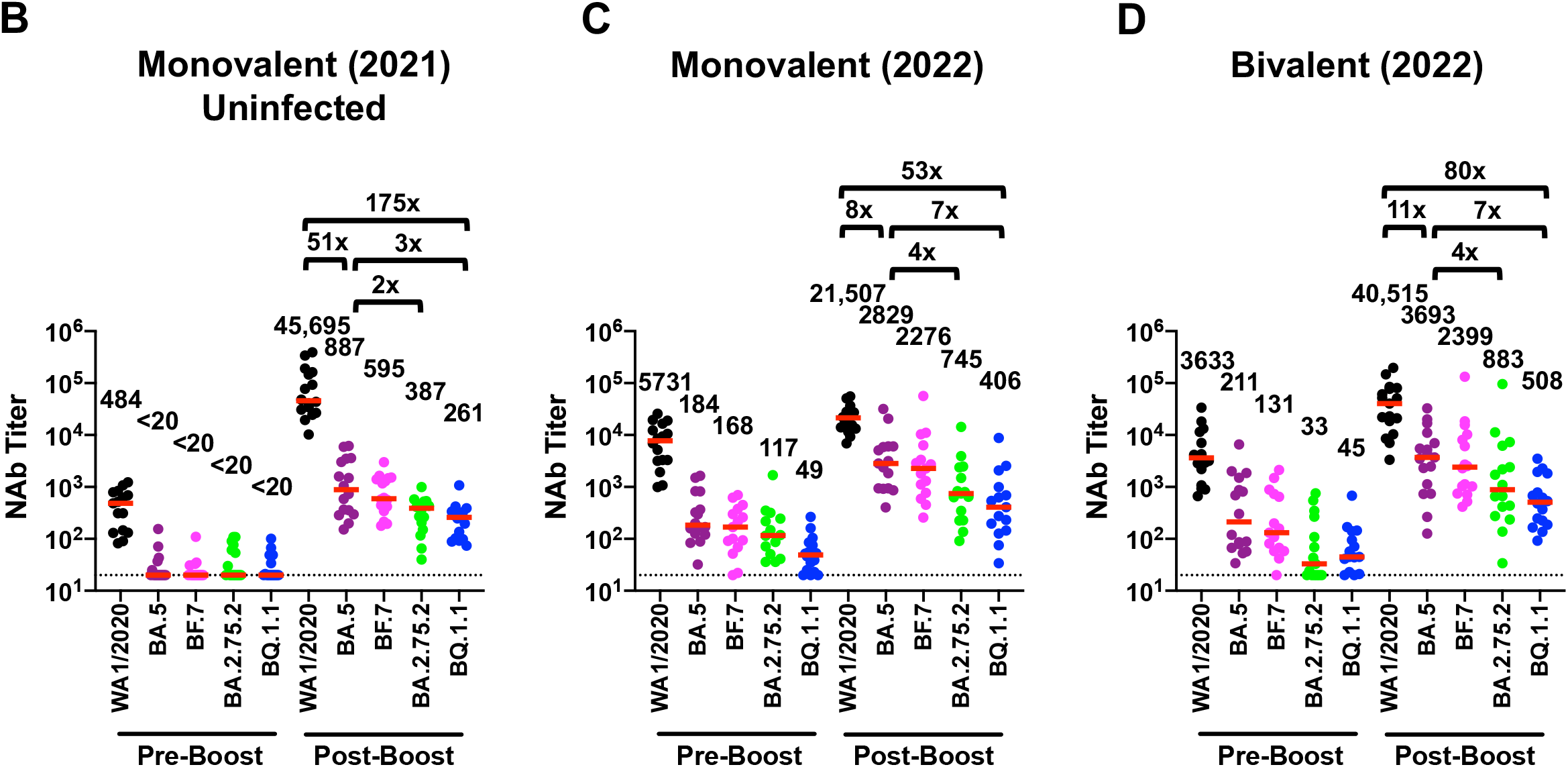
Neutralizing antibody responses to Omicron subvariants. **A**. Cartoon showing BA.1, BA.2, BA.2.75, BA.2.75.2, BA.5, BF.7, and BQ.1.1 mutations in the SARS-CoV-2 Spike. NTD, N-terminal domain; RBD, receptor binding domain; RBM, receptor binding motif; SD1, subdomain 1; SD2, subdomain 2; FP, fusion peptide; HR1, heptad repeat 1; HR2, heptad repeat 2. **B**. Neutralizing antibody (NAb) titers by a luciferase-based pseudovirus neutralization assay in uninfected individuals in 2021 six months following initial BNT162b2 vaccination (Pre-Boost) and following BNT162b2 boost (Post-Boost). **C**. NAb titers in individuals in 2022 before and after monovalent mRNA boosting. **D**. NAb titers in individuals in 2022 before and after bivalent mRNA boosting. NAb responses were measured against the SARS-CoV-2 WA1/2020, BA.5, BF.7, BA.2.75.2, and BQ.1.1 variants. The WA1/2020 and BA.5 titers in panels C and D are part of a separate manuscript and are included here for comparison. Medians (red bars) are depicted and shown numerically with fold differences.

We first assessed NAb titers in 16 individuals who were vaccinated and boosted with the monovalent mRNA BNT162b2 vaccine in 2021; participants were excluded from this group if they had known SARS-CoV-2 infection or a positive nucleocapsid serology (**Table S1**). Following the boost, median NAb titers to WA1/2020, BA.5, BF.7, BA.2.75.2, and BQ.1.1 were 45,695, 887, 595, 387, and 261, respectively (**Fig. 1B**). Median BQ.1.1 NAb titers were lower than median WA1/2020 and BA.5 NAb titers by factors of 175 and 3, respectively.

We next evaluated NAb titers in 15 individuals who received the monovalent mRNA boosters and in 18 individuals who received the bivalent mRNA boosters in 2022, most of whom received 3 (range 2-4) prior COVID-19 vaccine doses (**Table S1**). In these cohorts, 33% had documented SARS-CoV-2 Omicron infection, but we suspect that the majority were likely infected. Prior to boosting, WA1/2020 and Omicron NAb titers were higher in these groups (**Fig. 1C, D**) than in the uninfected 2021 cohort (**Fig. 1B**). Following the boost, median NAb titers to WA1/2020, BA.5, BF.7, BA.2.75.2, and BQ.1.1 were 21,507, 2829, 2276, 745, and 406, respectively, for the monovalent boosters (**Fig. 1C**) and were 40,515, 3693, 2399, 883, and 508, respectively, for the bivalent boosters (**Fig. 1D**). Median BQ.1.1 NAb titers were lower than median WA1/2020 and BA.5 NAb titers by factors of 53-80 and 7, respectively.

Our data demonstrate that BA.2.75.2 and BQ.1.1 escape NAbs induced by infection and vaccination more effectively than BA.5. BQ.1.1 NAb titers were lower than BA.5 NAb titers by a factor of 7 in two cohorts of individuals who received the monovalent or bivalent mRNA vaccine boosters. These findings provide the immunologic context for the rapid increase in BQ.1.1 prevalence in regions where BA.5 is dominant and have implications for both vaccine immunity and natural immunity. The incorporation of the R346T mutation into multiple new SARS-CoV-2 variants suggests convergent evolution.

## Data sharing

J.M., N.P.H., A.Y.C., and D.H.B. had full access to all the data in the study and take responsibility for the integrity of the data and the accuracy of the data analysis. All data are available in the manuscript or the supplementary material. Correspondence and requests for materials should be addressed to D.H.B.

## Supplementary Appendix

### Supplementary Methods

#### Study population

A specimen biorepository at Beth Israel Deaconess Medical Center (BIDMC) obtained samples from individuals who received SARS-CoV-2 vaccines as well as monovalent or bivalent mRNA boosters. The BIDMC institutional review board approved this study (2020P000361). All participants provided informed consent. This study included 16 uninfected individuals who received the original monovalent mRNA booster BNT162b2 in 2021. Participants were excluded from this group if they had a history of SARS-CoV-2 infection or a positive nucleocapsid (N) serology by electrochemiluminescence assays (ECLA), or if they received other COVID-19 vaccines or immunosuppressive medications. This study also included 15 individuals who received the original monovalent mRNA boosters BNT162b2 or mRNA-1273 between January and August 2022 and 18 individuals who received the bivalent mRNA boosters in September 2022 with various vaccination and infection history backgrounds. Participants were excluded from these groups if they received immunosuppressive medications.

#### Pseudovirus neutralizing antibody assay

Neutralizing antibody (NAb) titers against SARS-CoV-2 variants utilized pseudoviruses expressing a luciferase reporter gene. In brief, the packaging construct psPAX2 (AIDS Resource and Reagent Program), luciferase reporter plasmid pLenti-CMV Puro-Luc (Addgene), and Spike protein expressing pcDNA3.1-SARS-CoV-2 SΔCT were co-transfected into HEK293T cells (ATCC CRL_3216) with lipofectamine 2000 (ThermoFisher Scientific). Pseudoviruses of SARS-CoV-2 variants were generated using the Spike protein from WA1/2020 (Wuhan/WIV04/2019, GISAID ID: EPI_ISL_402124), Omicron BA.5 (GISAID ID: EPI_ISL_12268495.2), BF.7 (GISAID ID: EPI_ISL_15379594), BA.2.75.2 (GISAID ID: EPI_ISL_14913457), and BQ.1.1 (GISAID ID: EPI_ISL_14752457). The supernatants containing the pseudotype viruses were collected 48h after transfection, and pseudotype viruses were purified by filtration with 0.45-μm filter. To determine NAb titers in human serum, HEK293T-hACE2 cells were seeded in 96-well tissue culture plates at a density of 2 × 10^4^ cells per well overnight. Three-fold serial dilutions of heat-inactivated serum samples were prepared and mixed with 50 μl of pseudovirus. The mixture was incubated at 37 °C for 1 h before adding to HEK293T-hACE2 cells. After 48 h, cells were lysed in Steady-Glo Luciferase Assay (Promega) according to the manufacturer’s instructions. SARS-CoV-2 neutralization titers were defined as the sample dilution at which a 50% reduction (NT50) in relative light units was observed relative to the average of the virus control wells.

**Table S1.** Study Population.

|  | **Monovalent mRNA**<br>**Booster (2021)**<br>N=16 | **Bivalent mRNA**<br>**Booster (2022)**<br>N=18 | **Monovalent mRNA**<br>**Booster (2022)**<br>N=15 |
| --- | --- | --- | --- |
| **Age** (years), median (range) | 34 (23-62) | 42 (37-57) | 50 (33-64) |
| **Sex at birth**, Female | 15 (94) | 11 (61) | 13 (87) |
| **Race** |  |  |  |
| White | 12 (75) | 16 (89) | 10 (67) |
| Asian | 2 (13) | 2 (11) | 2 (13) |
| Black | 1 (6) | 0 | 2 (13) |
| More than one race | 1 (6) | 0 | 0 |
| Other | 0 | 0 | 1 (7)* |
| **Ethnicity** |  |  |  |
| Hispanic or Latino | 2 (13) | 0 | 1 (7) |
| Non-Hispanic | 14 (88) | 17 (94) | 14 (93) |
| Unknown | 0 | 1 (6) | 0 |
| **Medical condition** |  |  |  |
| Hypertension | 3 (19) | 2 (11) | 3 (20) |
| Diabetes | 0 | 1 (6) | 1 (7) |
| Pregnant | 2 (13) | 1 (6) | 2 (13) |
| Asthma | 0 | 2 (11) | 0 |
| **Most recent COVID-19 vaccine booster** |  |  |  |
| Pfizer monovalent booster | 16 (100) | N/A | 5 (33) |
| Moderna monovalent booster | N/A | N/A | 10 (67) |
| Pfizer bivalent booster | N/A | 8 (44) | N/A |
| Moderna bivalent booster | N/A | 10 (53) | N/A |
| **Prior COVID-19 vaccine history** |  |  |  |
| BNT (4 doses) | 0 | 2 (11) | 0 |
| BNT (3 doses) | 0 | 8 (44) | 2 (13) |
| BNT (2 doses) | 16 (100) | 0 | 0 |
| BNT / BNT / 1273 (3 doses) | 0 | 2 (11) | 1 (7) |
| BNT / BNT / Ad26 (3 doses) | 0 | 1 (6) | 4 (27) |
| 1273 (3 doses) | 0 | 3 (17) | 6 (40) |
| 1273 / BNT (2 doses) | 0 | 0 | 1 (7) |
| Ad26 / BNT / 1273 (3 doses) | 0 | 1 (6) | 0 |
| Ad26 / 1273 (2 doses) | 0 | 1 (6) | 1 (7) |
| **Days from last vaccine dose to sampling** | 20 (15-28) | 21 (16-23) | 32 (17-64) |
| **Known COVID-19 positive test** | 0 | 6 (33) | 5 (33) |
| **Days from positive test to last vaccine** | N/A | 166 (149-257)** | 143 (103-183)** |

BNT=BNT162b2 (Pfizer); 1273=mRNA-1273 (Moderna); Ad26=Ad26.COV2.S (Janssen)
Data displayed as median (range or interquartile range, IQR) and n (%);PCR, polymerase chain reaction; pregnant designation reflects time of last vaccine dose. All individuals with known prior infection had mild disease.
\*Self-reported race as Latina
\*\*Reported for only those with known prior infection

**Figure S1.**
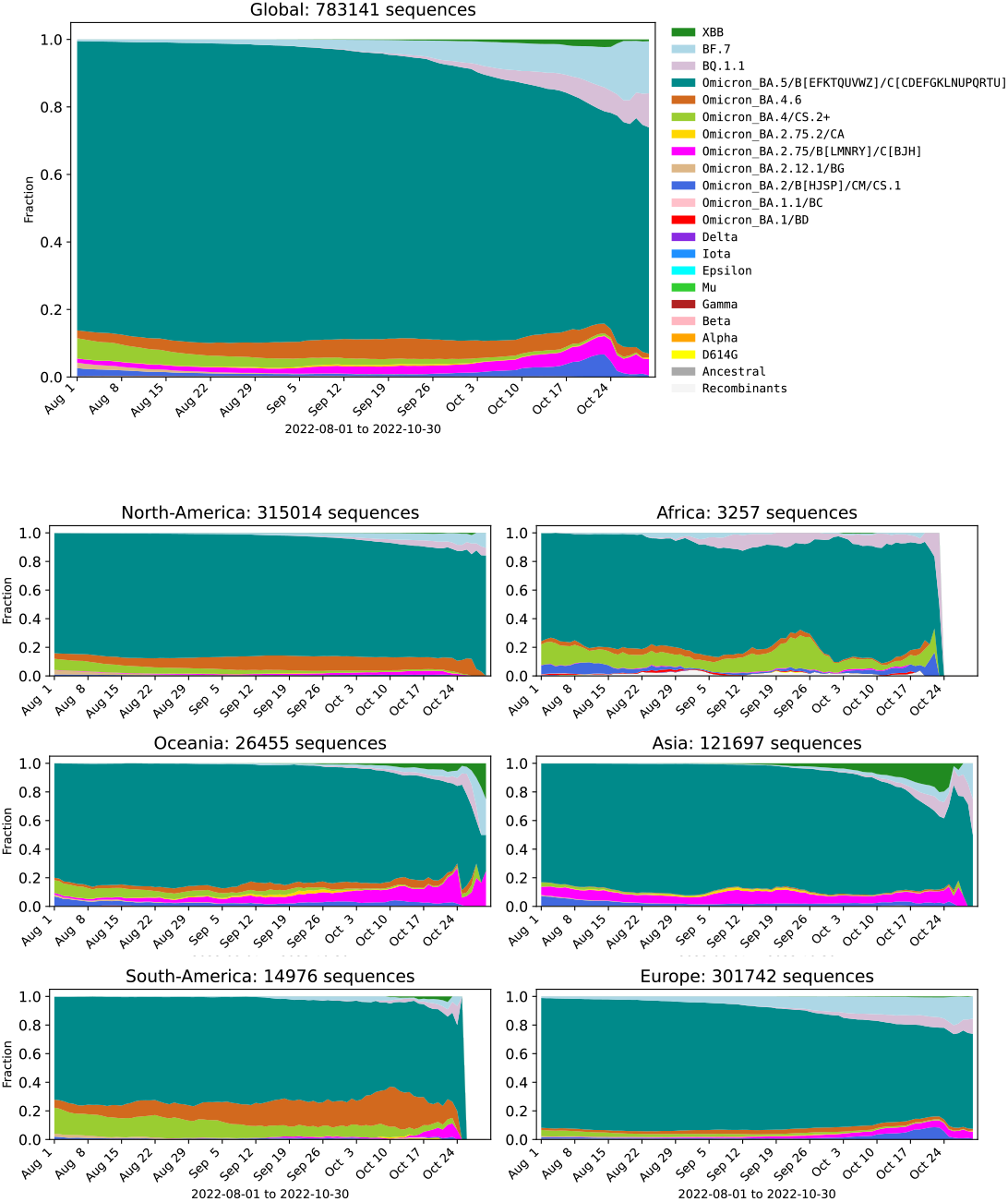
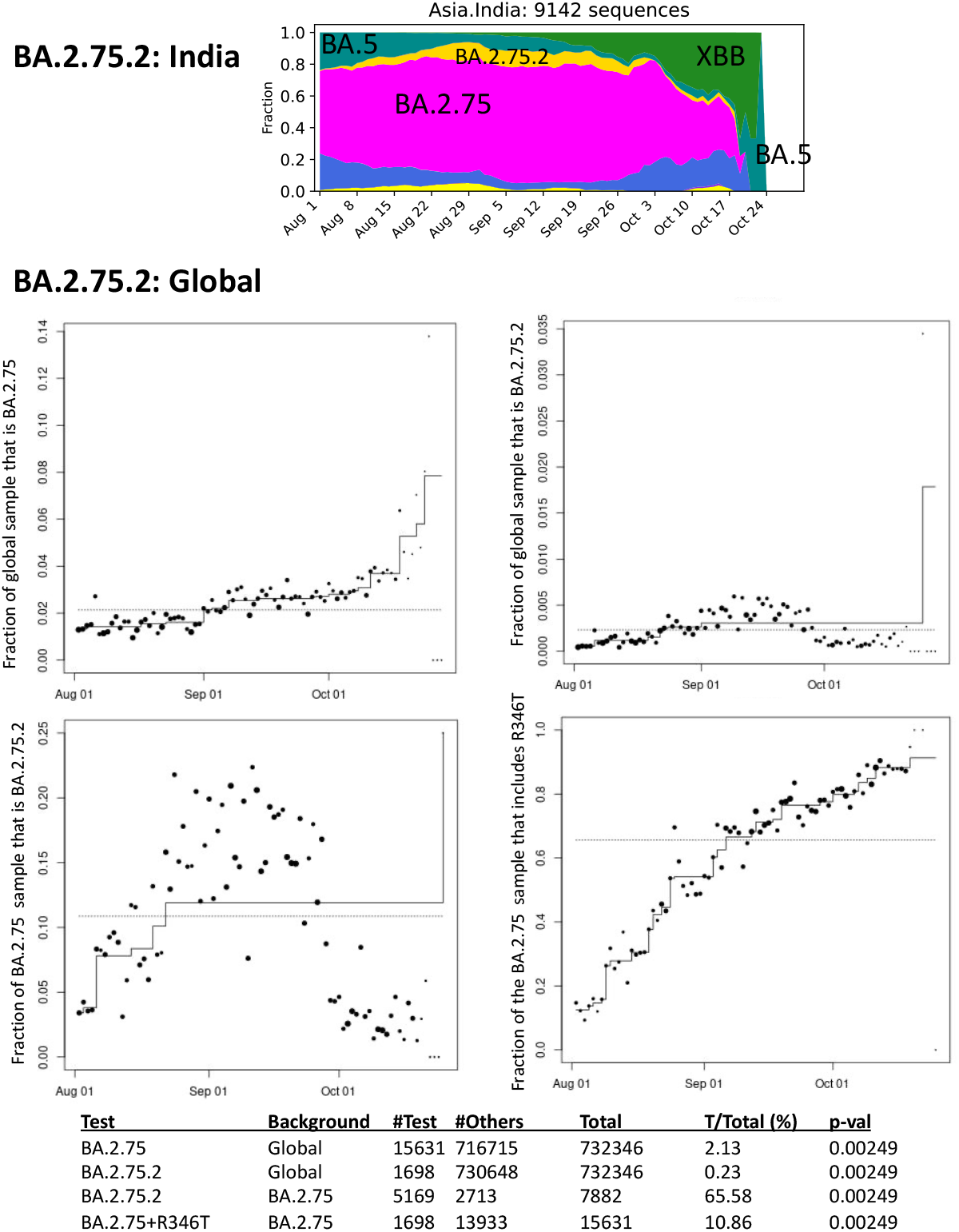
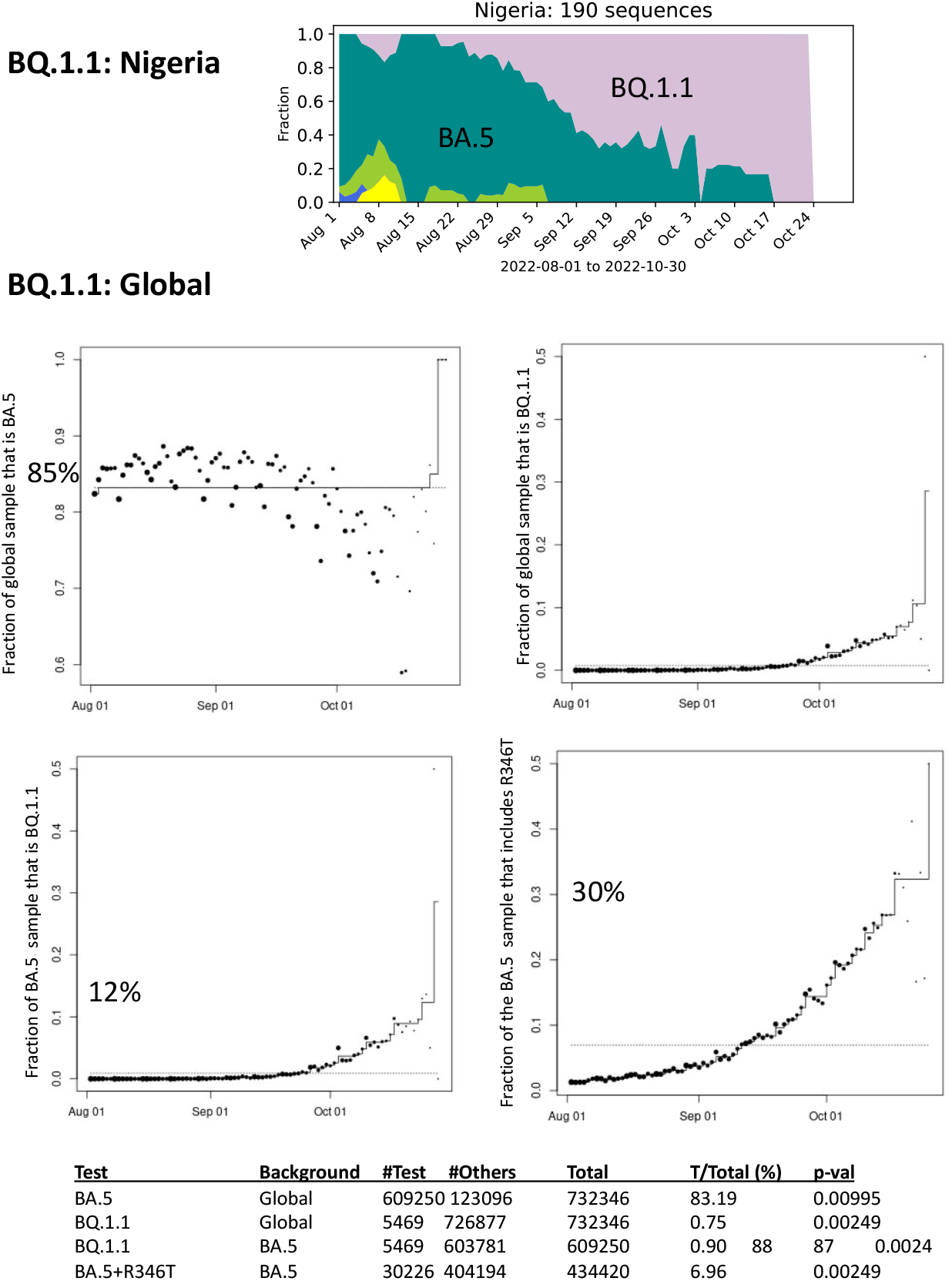
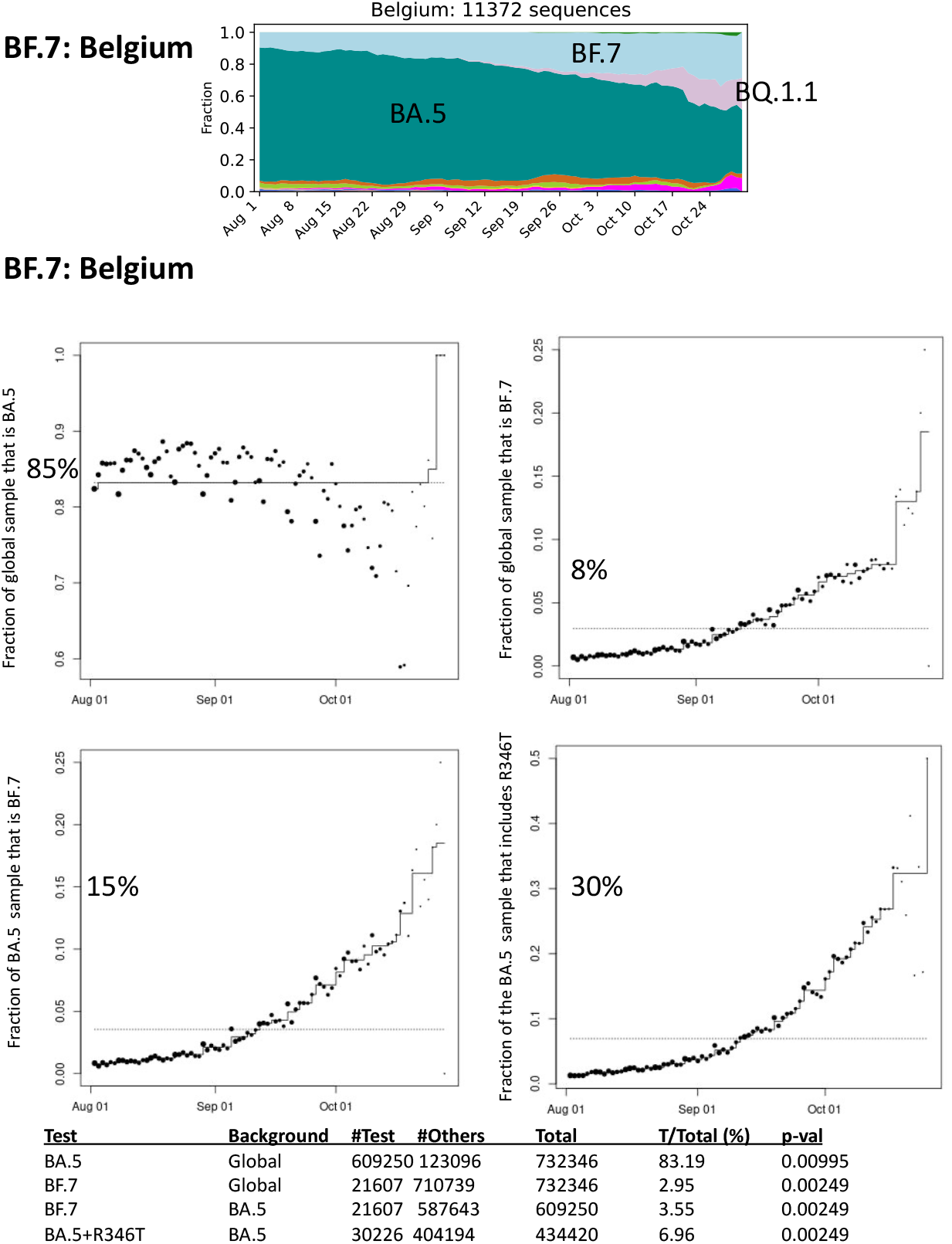
SARS-CoV2 Evolution. BA.2.75 has been gradually increasing globally but with only a modest increase in the BA.2.75.2 sublineage, although ∼90% of all BA.2.75 variants share its R346T mutation. BQ.1.1 has the R346T mutation and has displayed a particularly rapidly increase in frequency, suggesting that the K444T and N460K mutations may confer further selective advantage. BF.7 has the R346T mutation in the BA.5 sublineage but without these additional mutations.

## References

1. Barouch DH. Covid-19 Vaccines - Immunity, Variants, Boosters. N Engl J Me. 2022.

2. Hachmann NP. Miller J, Collier AY. et al. Neutralization Escape by SARS-CoV-2 Omicron Subvariants BA.2.12.1, BA.4, and BA.5. N Engl J Me. 2022;387:86–8.

3. Wang Q, Guo Y, Iketani S, et al. Antibody evasion by SARS-CoV-2 Omicron subvariants BA.2.12.1, BA.4 and BA.5. Nature 2022;608:603–8.

4. Hachmann NP. Miller J, Collier AY. Barouch DH. Neutralization Escape by SARS-CoV-2 Omicron Subvariant BA.4.6. N Engl J Me. 2022.

